# Universal enzyme-linked immunosorbent assays (ELISA) and utility in the detection of antibodies against Salmonella spp. in several animal species

**DOI:** 10.1101/2020.10.10.334771

**Authors:** Angel A Justiz-Vaillant, Belkis Ferrer-Cosme, Suzette Curtello

**Affiliations:** Department of Para-clinical Sciences. Faculty of medical Sciences. University of the West Indies. St. Augustine. Trinidad and Tobago. West Indies; Higher Institute of Medical Sciences of Santiago de Cuba. Cuba; Biotechnology Centre and Biochemistry Section of The Department of Basic Medical Sciences. University of the West Indies. Mona Campus. Jamaica. West Indies

**Keywords:** Immunoglobulin-binding bacterial protein (IBP), *Salmonella* spp, epidemiological survey

## Abstract

The aim of this study was to confirm the feasibility of using hybrid immunoglobulin-binding reagents in ELISAs for IgG/IgY detection and detection of specific antibodies against an infectious microorganism (*Salmonella* spp.) in various animal species using a universal diagnostic ELISA. Hybrid immunoglobulin-binding bacterial proteins (IBP), including recombinant protein LA, recombinant protein LG, and recombinant protein AG, have been produced to improve their binding affinity to a much larger number of immunoglobulins. Thus, this hybrid bacterial protein represents a powerful tool for the binding, detection, and purification of immunoglobulins and their fragments. However, SpLA-LG-peroxidase and SpLAG-anti-IgY-peroxidase were produced using the periodate method. These compounds have been shown to be effective as reagents. Their binding affinity to immunoglobulins surpasses that of previously reported hybrid IgG-binding proteins, including the most known SpAG, SpLA, and SpLG. The IgY fraction was isolated from the egg yolks of various birds, including chicken, bantam hen, guinea hen, quail, goose, duck, wild and domestic pigeons, parakeets, cattle egrets, pheasants, and ostrich. The IgY fraction was isolated using the chloroform-polyethylene glycol (PEG) method. An ELISA for anti-*Salmonella* spp. antibodies was employed with some modifications to determine the presence of antibodies in humans, laying hens, geese, quails, and pigeons. *Salmonella* is a motile, flagellated, rod-shaped zoonotic pathogen that can survive in the presence or absence of oxygen. They belong to the family Enterobacteriaceae and are implicated in typhoid fever and food-borne illnesses. This pathogen is associated with several diseases, which may become fatal and negatively impact the health of individuals and various economies globally. The poultry industry is most vulnerable to the influence of this pernicious microbe. The authors concluded that universal enzyme-linked immunosorbent assays were effective and reproducible in detecting immunoglobulins from both avian and mammalian species, but ELISA using the SpLAG-anti-IgY-HRP conjugate only reacted with the whole panel of animal antibodies. This conjugate was further used to standardize a universal ELISA for the determination of anti-Salmonella antibodies, in which human and avian species were serologically assessed.

## Introduction

Hybrid immunoglobulin-binding bacterial proteins (IBP), including recombinant protein LA, recombinant protein LG, and recombinant protein AG, have been produced to improve their binding affinity to a much larger number of immunoglobulins. In addition, they have been used in the diagnosis of zoonotic infections in tens of zoo animals worldwide. Thus, this hybrid bacterial protein represents a powerful tool for the binding, detection, and purification of immunoglobulins and their fragments [1].

By chemical protein conjugation, we engineered various hybrid bacterial Ig receptors with the capacity to bind to many immunoglobulins from mammalian and avian species. It includes SpLA-LG-peroxidase and SpLAG-anti-IgY-peroxidase produced by the periodate method [5].These molecules display a very low background in immunoassays. This makes them feasible as conjugates or coated to the solid phase of ELISAs for the detection of specific antibodies [2–4].

In a previous study, the binding affinities of individual immunoglobulin-binding reagents were tested, and researchers realized that none of the individual proteins, including protein-A, G, and L, bind extensively. None of these Ig-binding reagents bind with good binding affinity to immunoglobulins. However, their fusions, including protein AG (SpAG), protein LG SpLG) and protein LA (SpLA), were more effective Figure 1 shows SDS-PAGE, which depicts the creation of fusion proteins to detect antibodies. Several hybrid proteins, which have been extensively studied, can be visualized by protein electrophoresis [1].

**Figure 1.**
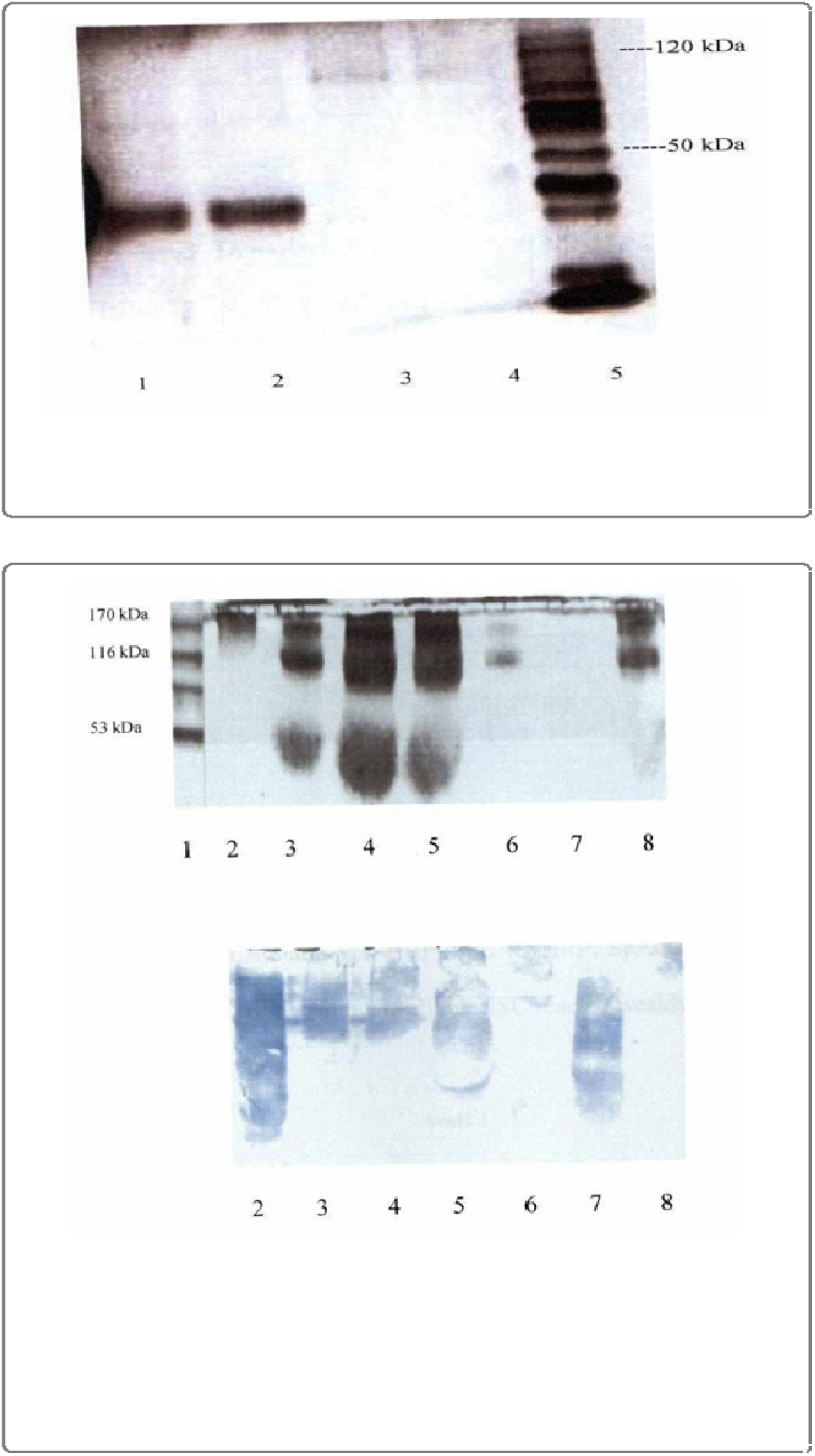
10% Denaturing SDS-PAGE of SpLA-LG-HRP conjugate in lanes 1-2 about 30 kDa and SpLAG-anti-IgY-HRP conjugate in lanes 3-4 that was denatured by heat and the protein bands are very faint. Lane 5 is the molecular weight (MW) marker. **Figure 2** shows denaturing SDS-PAGE and immunoblot analysis of chimeric conjugates: lane 1 MW marker, lane 2 SpLA-HRP (Sigma). The next conjugates were homemade lane 3 SpLAHRP, lane 4 SpLG-HRP), lane 5 SpG-HRP, lane 6 SpAG-HRP, lane 7 SpLA-LG-HRP and lane 8 SpLAG-HR. Immunoblot analysis shows that all chimeric conjugates interacted with human serum immunoglobulins and it confirmed their polymeric nature. Line 7 shows SpLA-LG-HRP a conjugate that will be used as a reagent in the development of a universal ELISA [1].

*Salmonella* are motile, flagellated, rod-shaped zoonotic pathogens that may survive with or without oxygen and do not absorb crystal violet stain. They are decolorized by alcohol because of their outer lipopolysaccharide membrane and thin peptidoglycan layer. They belong to the family Enterobacteriaceae and are implicated in typhoid fever and food-borne illnesses. This pathogen is associated with intestinal diseases that may become fatal and has a negative impact on the health of individuals and various economies globally [5]. The poultry industry is most vulnerable to the influence of this pernicious microbe. The lipopolysaccharide somatic O antigen, flagellar H, and virulent V antigenic structures determine the serotype designate *Salmonella* species [1].

Salmonellosis is an inflammation of the intestinal mucosal lining caused by *Salmonella* bacterial infiltration, resulting in painful symptoms. Both humans and animals are vulnerable. Salmonellosis is caused by the ingestion of *Salmonella* in food. The main route of transmission of *Salmonella* microbes is fecal-oral. Ingestion of small concentrations of *Salmonella,* such as six cells with contaminated food or water, will lead to salmonellosis infection. *Salmonella* resists the acidic conditions imposed by gastric juice and amino acids in the stomach. Incubation may last for six to twenty-four hours [1].

The bacteria adhere to receptors on the epithelial membrane of the intestine, where they stimulate enterotoxins, resulting in inflammation. One may experience stomach cramps, vomiting, nausea, diarrhea, ulcers, and elevated body temperature. Systemic breach of blood vessels and entry of bacteria into the bloodstream may be fatal [1]. Salmonellosis persists for up to 4-7 days. The disease usually resolves itself, and there is usually no need for antibiotics in healthy individuals, although it is commonly administered, especially among the elderly, immunocompromised individuals, and young children.

*Salmonella* enteritidis is a global problem with the most implicated cases of salmonellosis in Asia, the Americas, Africa, and Europe. *Salmonella* typhimurium presently leads to Enteritidis as a prominent source of salmonellosis in America. *Salmonella* Enteritidis serotype is associated with many deaths and gastroenteritis cases in the United States between 1990 and 2001. This incurred a loss of over eight hundred million dollars in the United States. There is concern that for every twenty thousand eggs produced, one will be contaminated by *Salmonella* Enteritidis, considering that there is production of more than 60 billion eggs annually by the industry [1].

The aim of this study was to confirm the feasibility of hybrid immunoglobulin-binding proteins used in ELISAs to study the binding affinity to IgG/IgY in many pet, laboratory, farm, and wild animals. We aimed to detect specific antibodies against infectious microorganisms (*Salmonella* spp.) in various animal species using a universal enzyme-linked immunosorbent assay (ELISA)6.

## Materials and Methods

### Samples for calculating the coefficient of variation (CV)

In this study, three samples of commercially available proteins (Sigma-Aldrich) were used along with a dolphin sample donated by the National Aquarium in Havana Cuba. Commercially purchased IgG samples of raccoons, skunks, and coyotes were purified as explained below and pooled to calculate the CV. Coefficients of variation (CV) were calculated using the formula = (standard deviation/mean) × 100. Human samples were purchased from Sigma-Aldrich.

### Immunoglobulins

The IgY fraction was isolated from the egg yolks of various birds, including chicken, bantam hen, guinea hen, quail, goose, duck, wild and domestic pigeons, parakeets, cattle egrets, pheasants, and ostrich. The IgY fraction was isolated using the chloroform-polyethylene glycol (PEG) method (Polson method, 1990) [7,8]. Purified immunoglobulin (Ig) was purified using an antibody purification system based on protein A-affinity chromatography (PURE-1A; Sigma-Aldrich). The manufacturer’s instructions were as described in [8]. Other mammalian IgGs used in this assay were purchased from Sigma-Aldrich or donated as described above.

### Conjugations of the immunoglobulin-binding bacterial antigens

Figure 1 and 2 show horseradish peroxidase (HRP) labelled SpA, SpG and/or SpL conjugates and their hybrid proteins as SpLAG-anti-IgY-HRP and SpLA-LG-HRP, which were prepared using the periodate method described by Nakane and Kawoi [9]. From all conjugate prepared SpLAGanti-IgY-HRP and SpLA-LG-HRP were less polymeric than other conjugates. The optimal working dilutions of the conjugates range from 1:2000 to 1:5000 [10].

**Figure 2.**
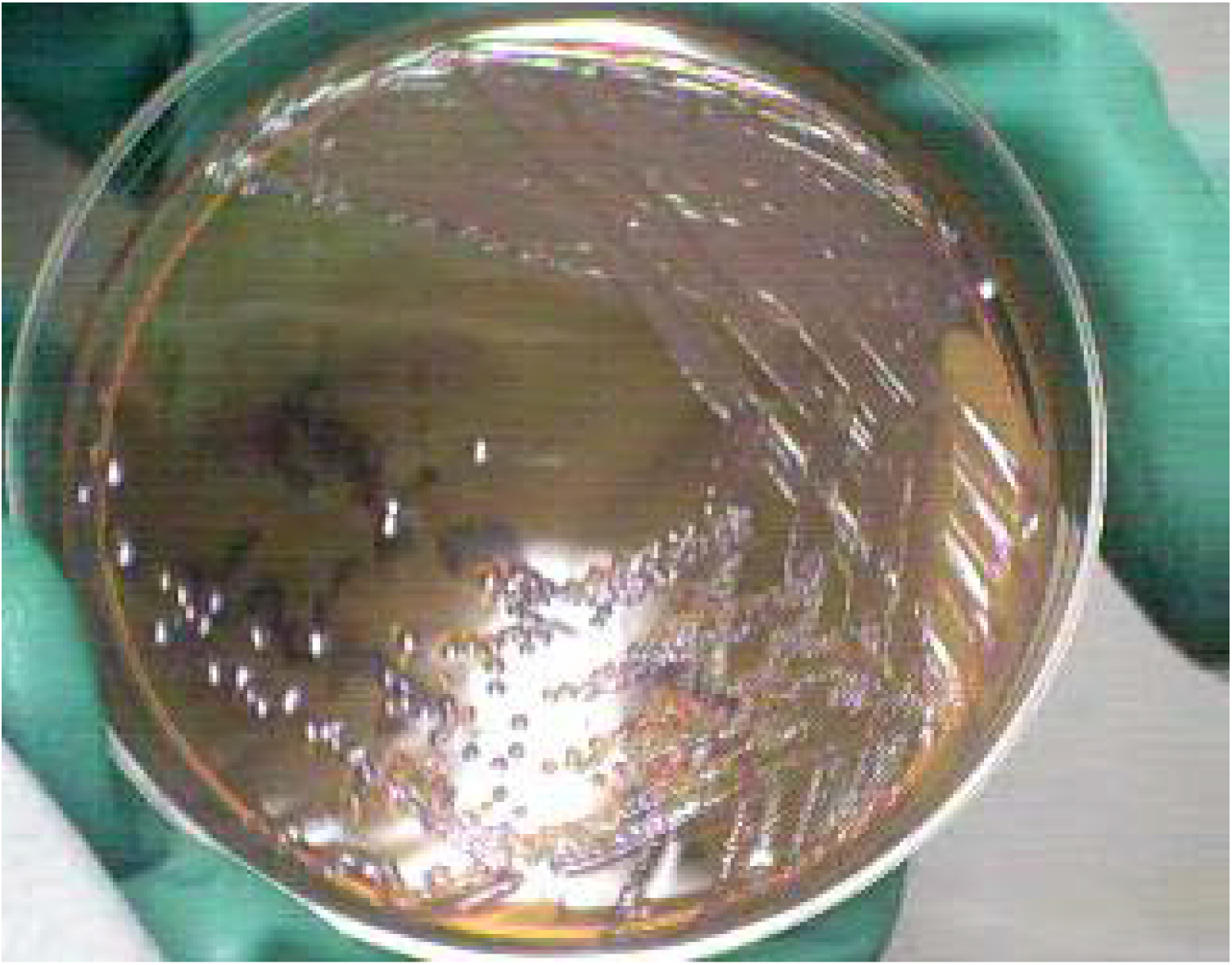
Typical isolated *Salmonella* colonies on differential culture media, MacConkey agar. Salmonellae are non-lactose fermenters and have a transparent appearance on MacConkey agar.

### Specimens

A total of 139 specimens were investigated for the presence of anti-Salmonella antibodies: eggs from laying hens (51), geese (10), quails (25) , and domestic pigeons (14). They were purchased at local grocery outlets, including supermarkets situated in the corporate areas of St. Andrew, in addition to local markets and a bird aviary where many eggs were collected. The egg specimens were aseptically placed in sterile bags. All egg samples were placed in an igloo with ice packs and transported to the laboratory [11–12]. Human samples were commercially available from Sigma-Aldrich Co.

The samples in question were IgYs from duck, ostrich, parakeet, Eagle egret, Bantam hen, pheasant, chicken, domestic pigeon, wild pigeon, goose, guinea hens, and quail. Mammalian IgGs, which bind to bacterial proteins, include antibodies from pigs, humans, skunks, coyotes, raccoons, mice, hamsters, bovines, goats, rabbits, cats, donkeys, mules, horses, dogs, pigs, and rats.

### Basic ELISA protocol using SpLA-LG-HRP and SpLAG-anti-IgY-HRP

The microplate was coated directly with avian or mammalian immunoglobulin in carbonate-bicarbonate buffer pH 9.6 overnight. The microwell was then treated with blocking solution and washed. Then, the bacterial protein conjugate was added and incubated for 1h. The microplate was rewashed and the enzyme substrate was added. After incubation in the dark for 15 min, the reaction was stopped, and the microwells were read at 450 nm. Positive samples were above the cut-off point, calculated from the mean of the negative controls.

### ELISA for anti-Salmonella antibodies in several animal species

An enzyme-linked immunosorbent assay was performed to detect anti-Salmonella antibodies in humans, and several avian species were detected. Ninety-six well polystyrene microplates (Ushaped bottom, Sigma-Aldrich) were incubated at 4°C overnight with 1 μg/well of LPS from Salmonella typhimurium. The microplates were washed and blocked with PBS-Tween 20 (pH7.4) for 1 h at room temperature (RT). The microplates were then washed four times as previously described. Then, 50 μL of human serum or 50 μL of IgY sample (1 mg/mL) from laying hens, geese, quails, and pigeons was added prior to IgY purification using the method of Polson [7]. After incubation for 1 h at RT, the microplates were washed again. Fifty (50) μl SpLAG-anti-IgY-HRP conjugate at a dilution of 1:5000 with PBS-Tween-20 was added. After a further incubation and washing procedure, 50μl tetramethylbenzidine (TMB) was added to each well. Microplates were further incubated for 15 min in the dark, and then 50μl of 3M HCl was added to each well to stop the reaction. microplates were read at 450 nm using a microplate reader. The cut-off value was calculated as mean optical density (XOD). The positive and negative controls (five each) were humans with very high or no titers of anti-Salmonella antibodies, respectively [1]. The intra- and inter-assay coefficients of variation were calculated as a measure of reproducibility.

### Statistical analysis

The Statistical Package for the Social Sciences (SPSS) version 25 was used to calculate the coefficients of variation and 95% confidence intervals. The proportions of antibodies and 95% confidence intervals were calculated for several animal species, including domestic pigeons, quails, and geese.

### Ethical approval

This study was approved by the Research Ethics Committee of the University of West Indies (UWI) (Mona Campus) in Jamaica, West Indies.

## Result and Discussion

### SpLA-LG Direct ELISA

This ELISA was used to study the interaction of SpLA-LG-HRP with avian and mammalian immunoglobulins. The 96 well microtiter plate was coated overnight at 4°C with 1 μg/μl per well of IgG or IgY (1 mg/mL) in carbonate-bicarbonate buffer (pH 9.6). The plate was then treated with bovine serum albumin solution and washed 4X with PBS-Tween. Then, 50 μL of peroxidase-labeled-SpLA-LG conjugate diluted 1:2000 in PBS-non-fat milk was added to each well and incubated for 1 h at RT. After that, the plate was washed 4X with PBS-Tween-20. Then, 50 μL of 3,3′,5,5′ - tetramethylbenzidine (TMB; Sigma-Aldrich) was pipetted onto each well. The reaction was stopped with 50 μL of a 3M H2SO4 solution. The plate was assessed visually for color development and read on a microplate reader at 450 nm. Subsequently, a cutoff point is calculated as the mean of the optical density of the negative controls multiplied by two [13]. The cut-off point was set to 0.268. Chicken IgY was used as a negative control, and affinity ranges were calculated.

Table 1 shows that SpLA-LG direct ELISA is a novel immunoassay that examines the binding affinity of a novel hybrid immunoglobulin-binding protein to immunoglobulins from a variety of animal species [10,13]. The authors are not aware of the development of an ELISA using this protein conjugate for immunodetection. The results showed that SpLA-LG preferentially binds to mammalian IgG. It binds strongly to 46.7% (14 out of 30 samples). It binds moderately to 13.33% of the immunoglobulin panel (4 out of 30 samples). SpLA-LG binds weakly to 10% of Igs (3 out of 30 samples) and does not bind to 30% of the Ig panel (9 out of 30). This protein binds specifically and preferentially to mammalian IgGs. However, it is not a good immunological marker for the detection of avian immunoglobulins, except for the IgY of bantam hen, ostrich, and duck. This hybrid protein was improved by the insertion of a molecule of antichicken IgY (Promega) in the SpLA-LG molecule, using the periodate method. Several ratios of bacterial protein-enzymes were assayed until the most appropriate ratio was 1:1. However, based on the SDS-PAGE results, an amount of unconjugated peroxidase was present in the conjugate, but it was eliminated with the washing steps after adding the labeled immunoglobulin-binding protein to the immunoassay.

**Table 1.** SpLA-LG Direct ELISA.

| **Immunoglobulins** | **XOD results** | **Binding affinity** | **Coefficient of variation (CV) intra-assay (%)** |
| --- | --- | --- | --- |
| Duck | 0.288 | + | 3.27 |
| Ostrich | 0.315 | + | 4.18 |
| Parakeet | 0.011 | - | 3.09 |
| Eagle egret | 0.090 | - | 3.02 |
| Bantam hen | 0.287 | + | 3.75 |
| Pheasant | 0.112 | - | 4.78 |
| *Chicken | 0.134 | - | 4.01 |
| Domestic pigeon | 0.089 | - | 2.89 |
| Wild pigeon | 0.083 | - | 3.11 |
| Goose | 0.108 | - | 4.05 |
| Guinea hen | 0.095 | - | 3.23 |
| Quail | 0.125 | - | 3.56 |
| Rat | 0.754 | ++ | 3.44 |
| Dog | 0.995 | +++ | 4.08 |
| Horse | 0.856 | +++ | 4.77 |
| Mule | 0.680 | ++ | 3.89 |
| Donkey | 0.814 | +++ | 3.70 |
| Cat | 0.655 | ++ | 3.68 |
| Rabbit | 1.058 | +++ | 4.18 |
| Goat | 0.886 | +++ | 5.17 |
| Bovine | 0.850 | +++ | 4.88 |
| Mouse | 1.122 | +++ | 4.22 |
| Raccoon | 0.907 | +++ | 4.06 |
| Coyote | 0.875 | +++ | 3.95 |
| Skunk | 1.259 | +++ | 4.05 |
| Hamster | 0.751 | ++ | 3.99 |
| Guinea pig | 1.004 | +++ | 4.72 |
| Human | 1.356 | +++ | 3.39 |
| Pig | 1.202 | +++ | 4.06 |
| Dolphin | 1.137 | +++ | 3.68 |

### Affinity ranges

XOD < 0.268= - (no binding)

0.268-0.535= + (weak)

0.536-0.803= ++ (moderate)

XOD > 0.804= +++ (strong)

### SpLAG-anti-IgY Direct ELISA

This ELISA was used to study the interaction of SpLAG-anti-IgY-HRP proteins with avian and mammalian immunoglobulins or antibodies [10,13]. The 96 well microtiter plate was coated overnight at 4°C with 1 μg/μl per well of IgG or IgY (1 mg/mL) in carbonate-bicarbonate buffer (pH 9.6). The plate was then treated with bovine serum albumin solution and washed 4X with PBS-Tween. Then, 50 μL of peroxidase-labeled-SpLAG conjugate diluted 1:2000 in PBS-nonfat milk was added to the microplate, which was then incubated for 1 h at RT. After that, the microplate was washed 4X with PBS-Tween-20. Then, 50 μL of 3,3′,5,5′ - tetramethylbenzidine (TMB; Sigma-Aldrich) was pipetted onto each well. The reaction was stopped with 50 μL of a 3M H2SO4 solution. The microwells were visually assessed for color development and read on a microplate reader at 450 nm. Subsequently, a cut-off point is calculated as the mean of the optical density of the negative controls multiplied by three [13,14]. The higher the OD value, the higher the binding affinity of the bacterial proteins to immunoglobulins. The cut-off point was set to 0.261.

The SpLAG-anti-IgY-HRP conjugate is one of the most versatile reagents for the standardization of universal ELISAs. It binds strongly to 45.16% of the Ig panel (14 out of 31 samples). It also binds moderately to 25.8% of immunoglobulins, including various avian and mammalian immunoglobulins as IgYs of ducks, bantam hens, ostriches, and chickens. It binds weakly to some immunoglobulin Y from avian species including parakeet, eagle egret, pheasant, both species of pigeons, goose, guinea hen, and quail. This conjugate surpasses the effectiveness of SpLA-LG-HRP, which partially binds to avian immunoglobulins. The SpLAG-anti-IgY-HRP with primary anti-chicken IgY antibodies that cross-react with all IgY of the panel gives advantages to the whole molecule, because it is the only conjugate with affinity for avian and mammalian antibodies.

The results of the binding affinity of bacterial proteins to immunoglobulins in the immunoassays suggest that neither the SpLA-LG-HRP nor the SpLAG-anti-IgY-HRP present steric hindrance in their interaction with antibodies, even when SpA and SpG bind to the Fc fragment of antibodies [14] and protein L binds to light chains or Fab regions of many antibodies [1]. It is more advantageous to use IBP than primary or secondary antibodies. In contrast, ELISAs that use peroxidase-labeled primary secondary antibodies are highly specific to animal species. For example, the detection of anti-Salmonella antibodies in humans using a specific human ELISA only allows the detection of antibodies in humans; however, when using any immunoglobulin-binding protein, combined with an enzyme, ELISA is a universal assay that allows the determination of specific antibodies in a greater number of species with a higher sensitivity and specificity. In the present study, anti-Salmonella antibodies were detected in five species, including a mammal (human) and four avian species, with only one standardized universal ELISA. Turtle IgY was used as a negative control because it does not bind to any immunoglobulin-binding protein [13,15].

**Table 2.** SpLAG-anti-IgY Direct ELISA [10,13].

| **Immunoglobulins** | **XOD results** | **Binding affinity** | **Coefficient of variation (CV) intra-assay (%)** |
| --- | --- | --- | --- |
| Duck | 0.513 | ++ | 2.84 |
| Ostrich | 0.550 | ++ | 3.89 |
| Parakeet | 0.364 | + | 4.74 |
| Eagle egret | 0.392 | + | 4.01 |
| Bantam hen | 0.580 | ++ | 3.65 |
| Pheasant | 0.329 | + | 4.13 |
| Chicken | 0.670 | ++ | 3.03 |
| Domestic pigeon | 0.387 | + | 3.59 |
| Wild pigeon | 0.305 | + | 4.52 |
| Goose | 0.315 | + | 4.38 |
| Guinea hen | 0.381 | + | 3.12 |
| Quail | 0.396 | + | 4.59 |
| Rat | 0.754 | ++ | 2.95 |
| Dog | 0.995 | +++ | 3.66 |
| Horse | 0.856 | +++ | 5.24 |
| Mule | 0.680 | ++ | 4.96 |
| Donkey | 0.814 | +++ | 4.62 |
| Cat | 0.655 | ++ | 3.26 |
| Rabbit | 1.058 | +++ | 3.56 |
| Goat | 0.886 | +++ | 4.64 |
| Bovine | 0.850 | +++ | 4.06 |
| Mouse | 1.122 | +++ | 3.50 |
| Raccoon | 0.907 | +++ | 3.44 |
| Coyote | 0.875 | +++ | 4.17 |
| Skunk | 1.259 | +++ | 3.61 |
| Hamster | 0.751 | ++ | 5.13 |
| Guinea pig | 1.004 | +++ | 3.57 |
| Human | 1.356 | +++ | 4.26 |
| Pig | 1.202 | +++ | 3.98 |
| Dolphin | 0.991 | +++ | 4.21 |
| Turtle IgY | 0.090 | - | 2.88 |

### Affinity ranges

XOD < 0.27= - (no binding)

0.27-0.53= + (weak)

0.54-0.81= ++ (moderate)

XOD > 0.81= +++ (strong)

Table 3 shows that laying hens (74.5%) and humans (22.5%) had the highest seroprevalence of anti-Salmonella antibodies among all species. ELISA using the SpLAG-anti-IgY-HRP conjugate was effective in the determination of both avian and mammalian immunoglobulins. Table 4 shows the different 95% confidence intervals, where the sample means fall in between. From 23.13-32.87 for laying hens, between 22.46-33.54 for humans, between 20.85-35.15 for quails, between 17.99 and 38.01 for domestic pigeons and between 15.6 and 40.4 for geese.

**Table 3.** Presence of anti-Salmonella antibodies in egg yolk and human serum by SpLAG-anti-IgY universal ELISA.

| Species | Positive (%) |
| --- | --- |
| Human | 22.5% (9/40) |
| Geese | 10.0% (1/10) |
| Quail | 16.0% (4/25) |
| Domestic Pigeon | 14.3% (2/14) |
| Laying hen | 74.5% (38/51) |

**Table 4.**
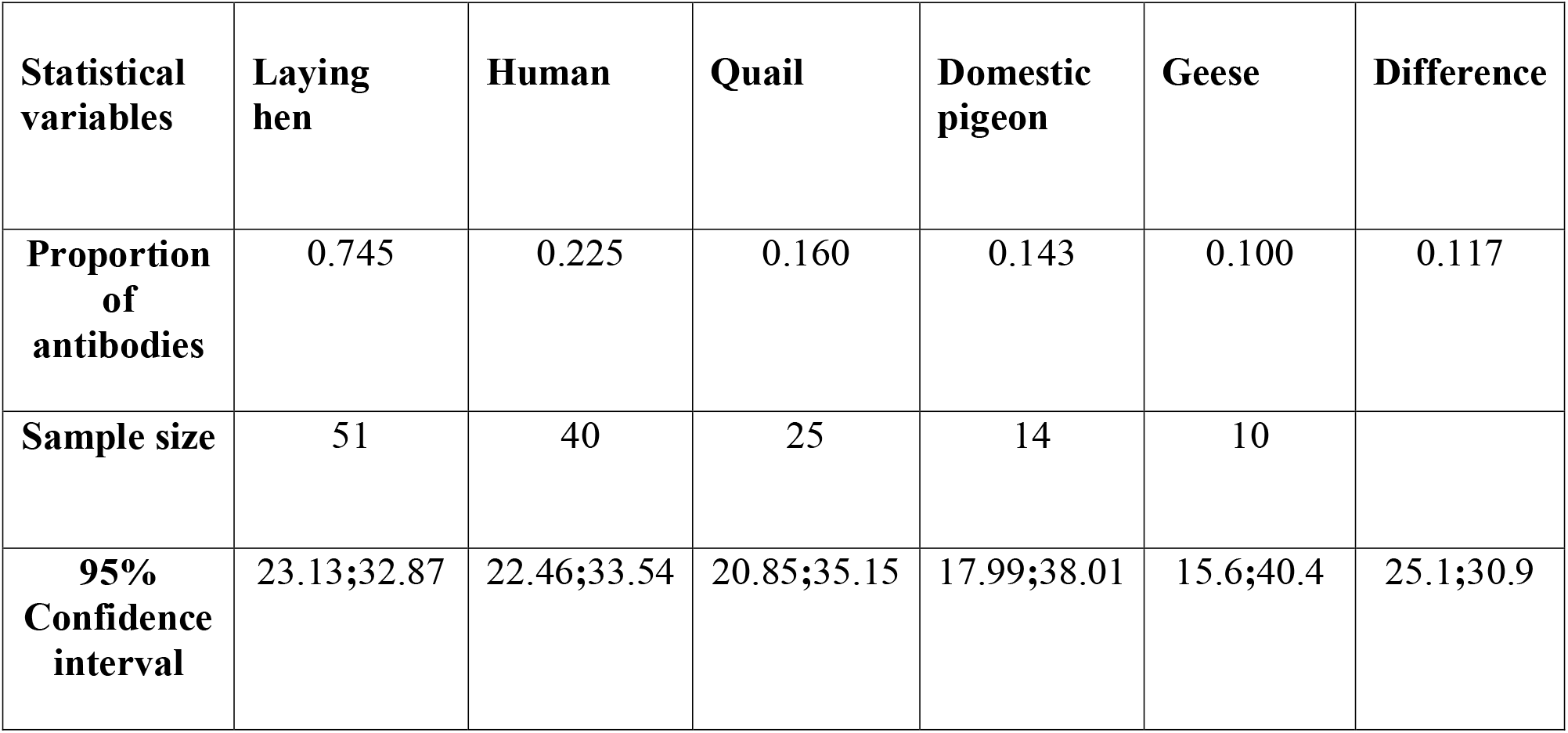
Table showing the differences in proportions between human and avian samples with 95% confidence intervals.

Table 4 shows The 95% confidence interval is a range of values that you can be 95% certain contains the true mean of the population as seen in commercial samples of humans (Sigma-Aldrich) and other animal species as laying hens, quails, geese, domestic pigeon and quails.

In a previous epidemiological study of Salmonellosis affecting humans and chickens, 11.3% of humans (6/53) and 95.3% of chicken (102/107) depicted anti-*Salmonella* antibodies in human sera and chicken egg yolks. *Salmonella* is ubiquitous, and chickens may be asymptomatic, which has generated great health concerns [12]. This bacterium is associated with life-threatening diseases in humans and animals. Gastroenteritis is a disease that may resolve within five days in healthy individuals. Immunocompromised individuals and very young individuals are at great risk, as this may progress to secondary systemic complications [1].

The frequency and ease of *Salmonella* enteritidis infection in humans due to their close association with eggs make it vulnerable to the spread of salmonellosis globally. The World Health Organization 2001 data labeled *Salmonella* Enteritidis as a global problem with the most implicated cases of salmonellosis in Asia, the Americas, Africa, and Europe. *Salmonella* typhimurium presently leads to Enteritidis as a prominent source of salmonellosis in America. *Salmonella* Enteritidis serotype is associated with many deaths and gastroenteritis cases in the United States between 1990 and 2001. This incurred a loss of over eight hundred million dollars in the United States. There is concern that for every twenty thousand eggs produced, one will be contaminated by *Salmonella* Enteritidis, considering that there is production of more than 60 billion eggs annually by the industry [16].

The mystery surrounding the capacity of *Salmonella* Enteritidis to infiltrate and taint intact eggs has led to the hypothesis that this pathogen has the ability to inhabit the ovary and oviduct of layer chickens infiltrating the eggs as they are formed. Other *Salmonella* serotypes, such as Typhimurium, only gain access to eggs when contaminated chicken crosses contaminate cracked eggs. The resonating question by Keller et al. (1995) has been, why has *Salmonella* Enteritidis, overwhelmed industrialized countries for the past thirty years, with a penchant for egg products [16]. Callaway et al. (2008) noted that the elimination of the serotype Gallinarum linked to chicken typhoid in the early twentieth century may have led to the environmental and ecological voids that have been exploited by *Salmonella* Enteritidis. Since the elimination of the serotype Gallinarum, subsequent years have steadily shown an increase in salmonellosis linked to the consumption of poultry products. The escalated insurgence of salmonellosis associated with poultry products has led to an overwhelming fear and stigmatization of uncooked poultry products as poison [17].

The cost to the United States economy in 2008 exceeded two billion dollars. Over the United States of America, there are five hundred deaths annually. *Salmonella* may adapt to various hosts, without any manifestation of infection. U.S. Food Safety surveillance plans to reduce *Salmonella* incidence to below 19 percent [17]. Vaccine implementation has been embraced in collaboration with Intervet and Southeast Poultry to improve existing vaccines against salmonellosis. Southeast vaccine designs outperformed others during trials; hence, collaboration to combine resources and expertise to ultimately develop an ideal vaccine with a protective immunological response [18].

Regulatory bodies and control processes are required to prevent and effectively prevent the spread of *Salmonella* infection and minimize cross-contamination. Close scrutiny and control by regulatory bodies of meat and poultry plants to promote safe food products is vital for combating salmonellosis. The implementation of a modern approach by the Agriculture Department of Poultry and Meat Inspection Program in the United States is an initial step in the prevention process. There is a requirement for the adoption of a system that controls the process of operation by slaughtering and processing plants for safe operations and product quality. Hazard analysis and critical control points (HACCP) are control systems that are introduced to eliminate hazards from food and maintain a safe food chain. The Food Safety and Reduction Service has implemented measures to verify the effectiveness of the HACCP Hazard system in the reduction of pathogens such as *Salmonella*. Abattoirs are active players in the drive to reduce harmful pathogens and maintain performance standards at all points of operation, both within the slaughter, processing plants, and farm locals.

Chicken feeds, rats, free-flying birds, flies, poultry house environment, processing plant environment, and transportation are some of the possible contamination sources that make control of *Salmonella* on a poultry farm difficult [19]. Vaccination prevents Salmonella colonization. The perceived view of the Department of Agriculture of the United States is that radiation will control *Salmonella*; however, this view is not endorsed by the majority of stakeholders. The incidence of salmonellosis may be controlled by synchronized measures applied to the different phases of production, processing, distribution retail marketing, and handling in the prevention and curtail of the propagation of *Salmonella*. Control measures must be supported by effective surveillance programs and the endorsement of all involved in its implementation [19, 20].

Consumers should be protected at all costs due to the exposure to *Salmonella* by tainted food. Careful handling of food that is not subjected to heat treatment or pasteurization before consumption must be practiced. Raw and uncooked food should be handled with the strictest of care to eliminate the possibility of cross-contamination. Appropriate refrigeration is practiced, and careful attention must be paid to the expiratory dates and proper cooking of foods. There should be a general separation of vegetables and fruits from raw poultry and meat. Regular hand washing should be practiced when handling meat and other food products. Raw materials and supplies for manufacturing should be sourced from suppliers who endorse and adhere to strict food safety hazard control measures.

Hygiene training for workers involved in food handling, processing plants, and critical areas of production within the plant and packaging areas There should be general hygiene training for all-inclusive farm workers, as the program for safe food should have a comprehensive inclusion approach. Consumers and operators of restaurants must be onboard and be cognizant of safe food practices, their role in keeping themselves, the environment, and ultimately the food chain free and void of *Salmonella*, the health hazards it presents to the general public.

## Conclusion

Universal enzyme-linked immunosorbent assays were effective and reproducible in detecting immunoglobulins from both avian and mammalian species, but ELISA using the SpLAG-anti-IgY-HRP conjugate only reacted with the whole panel of animal antibodies. This conjugate was further used to standardize a universal ELISA for the determination of anti-Salmonella antibodies, in which human and avian species were serologically assessed.

